# Microplastics analytics: why we should not underestimate the importance of blank controls

**DOI:** 10.1101/2023.02.04.527118

**Authors:** Michael J. Noonan, Nicole G. Ribeiro, C Lauren Mills, Marcia de A. M. M. Ferraz

## Abstract

Recent years have seen considerable scientific attention devoted towards documenting the presence of microplastics (MPs) in environmental samples. Due to omnipresence of environmental microplastics, however, disentangling environmental MPs from sample contamination is a challenge. Hence, the environmental (collection site and laboratory) microplastics contamination of samples during processing is a reality that we must address, in order to generate reproducible and reliable data. Here we investigated published literature and have found that around 1/5 of studies failed to use blank controls in their experiments. Additionally, only 34% of the studies used a controlled air environment for their samples processing (laminar flow, fume hood, closed laboratory, clean room, etc.). In that regard, we have also shown that preparing samples in the fume hood, leads to more microplastics contamination than preparing it in the laboratory bench and the laminar flow. Although it did not completely prevent microplastics contamination, the processing of sample inside the laminar flow is the best option to reduce sample contamination during processing. Overall, we showed that blank controls are a must in microplastics sample preparation, but it is often overlooked by researchers.

**Highlights:**

1. Most of the contaminant microplastics in blank controls were particles < 20 μm.
2. Fume hoods result in more contamination than processing the samples on the bench.
3. Laminar flow was the best option for reducing MPs contamination of samples.
4. 1/5 of studies failed to use blank controls, and 1/3 did not correct their data.
5. Improving the use and description of blanks is imperative for ensuring data quality.

**Mandatory graphical abstract:** 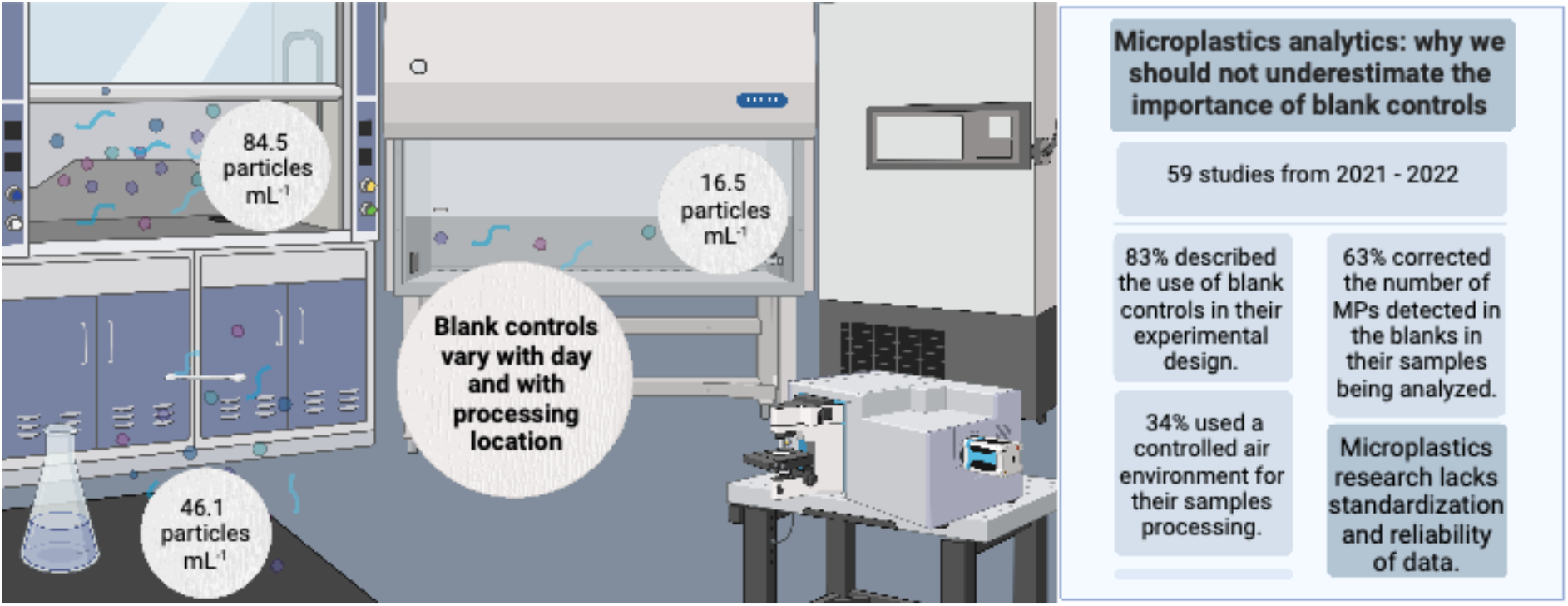

## 1. Introduction

Recent years have seen considerable scientific, public, and governmental attention devoted towards the plastic pollution crisis (European Commission, 2018; United Nations Environment Assembly, 2022), and studies on microplastics (MPs) are accumulating exponentially. The increasing number of peer-reviewed studies would suggest that studying MPs is a straightforward process. Nevertheless, a key challenge to meaningfully studying environmental MPs involves the fact that plastics are now so ubiquitous that contamination throughout the sample collection and/or processing steps is a nearly unavoidable reality. The high, and persistent risk of sample contamination makes it challenging to disentangle the actual presence of MPs in a sample from contamination, and the proper use of blank controls is thus essential for generating trustworthy data (Prata et al., 2020; Rillig et al., 2022; Tsering et al., 2022). As Rillig et al. (2022) point out, “*Controls are an essential component of experimental design, serving the purpose of accounting for all aspects of an experimental treatment except for the factor(s) under investigation in any given study*”. Yet, despite their critical importance, it is not uncommon to see MP studies without any controls. Thus, while the number of “first time reported” studies is growing (e.g., Hernandez et al., 2019; Li et al., 2020; Ragusa et al., 2021), the extent to which sample contamination may be playing a role in the discovery of plastics in different environmental and biological compartments is unknown.

Our aims in this study were twofold. First, to demonstrate the importance of controls, we prepared blank samples for MPs evaluation under three different laboratorial conditions: i) on the bench, ii) inside a fume hood and iii) inside a laminar flow – three of the most common ways of preparing blank controls in MPs research (e.g., Gardon et al., 2021; Paler et al., 2021; Xu et al., 2021). We then quantified the amount of MP contamination in each of these samples throughout an otherwise normal MP isolation and quantification process. As a further demonstration of the importance of blanks, we corrected a human follicular fluid sample that was previously analyzed in our laboratory (Grechi et al., 2022) to each of these blanks, and compared the results obtained under the different protocols. Second, we reviewed the open access published literature on MPs, to quantify the proportion of researchers that implemented blank controls in their experimental design(s), presented such data in their manuscripts, and corrected their results to the blank. The results of this work will provide important context to the growing volume of “first time reported” studies, and help guide future experimental work.

## 2. Material and methods

### 2.1. Experimental evaluation of blank control protocols

#### 2.1.1. MP contamination prevention

To reduce sample contamination with laboratory MPs, all flasks and other apparatuses were replaced by glass materials whenever possible. Moreover, all materials and equipment used were rinsed three times with filtered (0.1 μm filter – Merck Isopore) ultra-pure water prior to use. All reagents and water used in the protocols described below were also filtered using a 0.1 μm filter before use. All the materials washing and reagents filtering were performed inside a laminar flow. All vials were covered with clean aluminum foil during storage and incubations.

#### 2.1.2. Isolation of microplastics from water processed in the bench, fume hood or laminar flow

For all procedures, the same filtered ultra-pure water and other reagents were used. To prepare each group, 2 mL of filtered water were first placed in a pre-cleaned Erlenmeyer inside the laminar flow and covered with aluminum foil. From here on, each water sample was processed either on the bench, inside a fume hood or inside the laminar flow, at the same time. They were kept closed with the aluminum foil during all procedures, which was only removed when adding new solution or for filtration. In order to mimic the timeline and processes that the samples would normally go through, we used a protocol that was optimized for isolating and quantifying small microplastics from follicular fluid (Grechi et al., 2022). Briefly, on day one, KOH 10% in a proportion of 1:25 (sample : digestion solution) was added to each 2 mL of water, and incubated in a shaker at 250 rpm and 60ºC. After 24 h, NaClO was added to each Erlenmeyer to reach a final concentration of 0.84%, samples were incubated for 24 h at 250 rpm and 60ºC. All samples were then filtered in a 47 mm polytetrafluoroethylene polymer (PTFE) membrane (0.45 μm pores, Merck Millipore, USA) and rinsed three times with filtered ultra-pure water. The membranes were placed in a beaker containing 50 mL of HNO_3_ 20% and incubated in an ultrasonic bath (TI-H-5 MF2 230 V, Elma Schmidbauer GmbH, Germany) with 100% power, sweep function and a frequency of 45 kHz for 15 min (to transfer particles from the membrane into the solution), samples were then incubated at 40ºC and 250 rpm for another 24 h, filtered using 13 mm PTFE membranes (0.45 μm pores) and rinsed three times with ultra-pure filtered water. The membranes were mounted on a glass slide, left to dry at room temperature and used for microscopy and spectroscopy analysis. Samples were kept in a pre-cleaned glass petri dish until analysis to prevent further contamination.

#### 2.1.3. Analysis of isolated microplastics by confocal Raman spectroscopy

Following Grechi et al. (2022), a confocal Raman spectroscope (alpha300 R, WITec, Germany) with a spectrometer (UHTS 300, WITec, Germany), and a 532 nm laser was used for the characterization of the microplastics. The microscope was operated by using the Control Five software, which allowed the imaging of the whole membrane area using a ×50 NA 0.75 Zeiss objective. We then used the software Particle Scout to identify all the particles present in the membrane. The Raman spectra of each particle was acquired with a ×100 DIC NA 0.9 Zeiss objective, using the autofocus function, with a laser power of 10 mV, an accumulation of 4 and integration time of 0.45 s. The presence of microplastics was confirmed by matching the spectra found using the True Match Integrated Raman Spectra Database Management software, with the S.T. Japan database (S.T. Japan Europe GmbH, Germany) and the SloPP MPs Raman library (Munno et al., 2020). Only matches with a hit quality index (HQI) value higher than 75% were considered for further analysis. Poly(tetrafluoroethylene-co-perfluoro-(alkyl vinyl ether)) – PTFE – and Perfluoroalkoxy alkane particles were excluded from the analysis, since they were components of the pore membrane used for imaging. The particles were classified as: 1) non-plastic related particles; 2) plastic polymers; 3) plasticizers; 4) pigments; 5) coatings, solvents or fillers; 6) fiber; and 7) unknown. Data S1 provides a list of all different particles and their classification.

### 2.2. Analysis of the use of blank controls in the peer-reviewed literature

In order to select manuscripts to be included in our review, we performed a Pubmed (https://pubmed.ncbi.nlm.nih.gov/) search of the term “microplastic” on 15 August 2022. We then selected Open Access manuscripts that were published in the past 1 year (2021 – 2022) and the list of manuscripts was downloaded in .csv file format. The `R` function `sample()` was used to randomly sort the manuscripts and minimize any bias in the selection process. For the first 59 manuscripts, excluding review papers and manuscripts that did not quantify microplastics, we compiled data on: sample type, particle size, use of hood for sample processing/analyses, method of microplastics analysis, if a blank was used, if blank results were shown, what were the concentrations and sizes of microplastics in the blank, and if the samples were adjusted to the blank control. In order to ensure data were recorded correctly, the values from a random sample of 10 papers were confirmed by two co-authors. No discrepancies were noticed in this process. The PRISMA checklist is provided in Appendix S1 and the review process was not registered (Page et al., 2021).

## 3. Results and Discussion

### 3.1 Laboratory MPs contamination is a reality that needs to be tackled

To demonstrate the importance of the laboratory environment as a source of MPs contamination, we prepared and analysed water samples in a laminar flow cabinet, in a fume hood, and directly on a bench, which are the most common laboratory setups used to process samples in MPs studies (see Data S2). A large number of particles were identified in all the sample membranes, ranging from 1,975 to 27,329 total particles. Despite the large number of particles, only about 15% had Raman spectra with hit quality indices (HQIs) ≥ 75. Although there were differences in the number of particles for the three different types of blanks, the size profiles of the MP contaminants were relatively consistent (Fig. 1 a-c). The majority of the MP particles detected were smaller than 20 μm in length (90, 82 and 87.5 % of total MPs, for laminar flow, bench and fume hood samples, respectively), and only a small fraction of the MPs were larger than 100 μm in length (0, 0.3 and 0.4 % of total MPs, for laminar flow, bench and fume hood samples, respectively).

**Figure 1.**
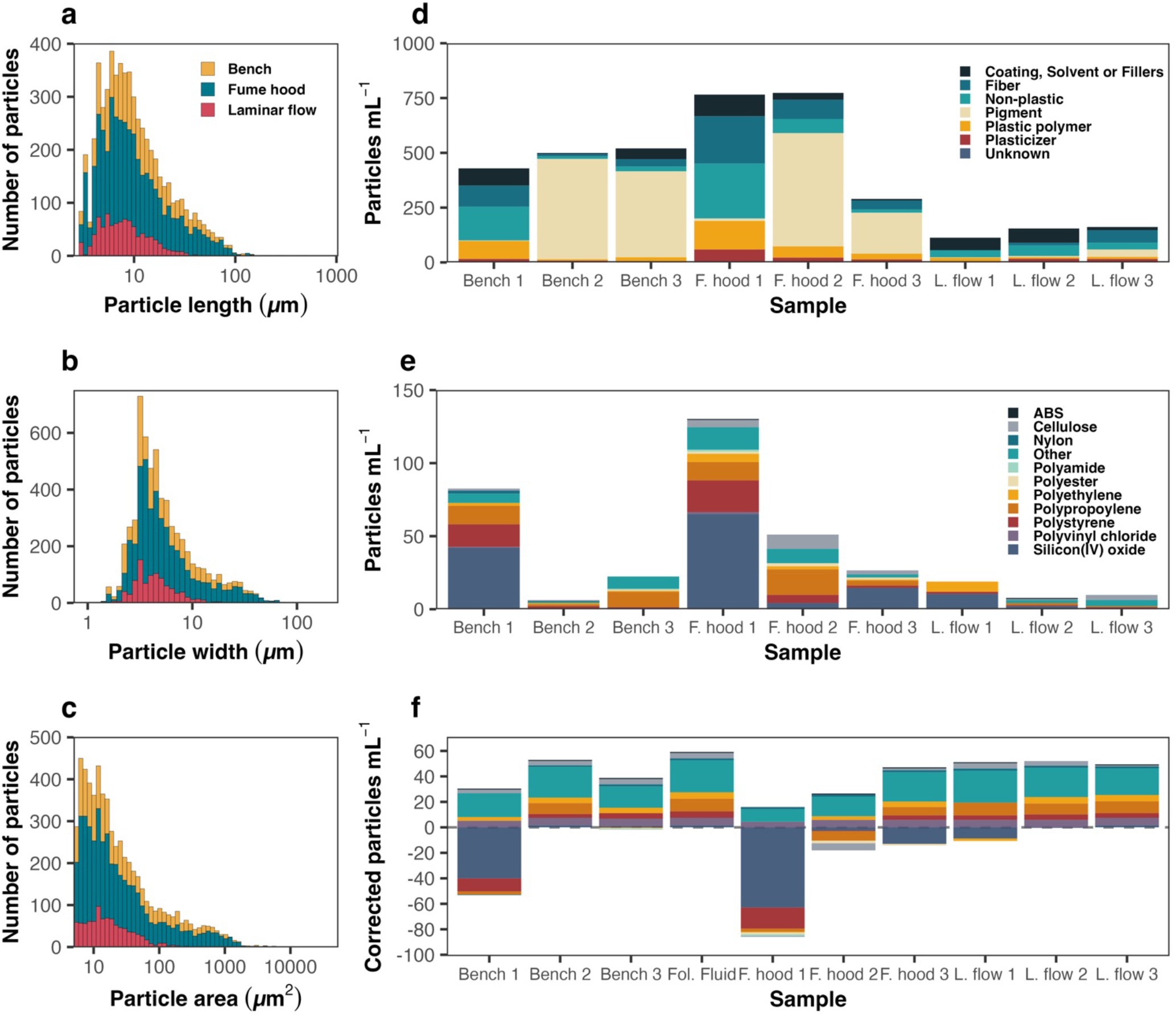
Characterization of blank control samples processed in a laminar flow cabinet (L. flow), a fume hood (F. hood), or directly on a bench (Bench) by confocal Raman spectroscopy. Histograms of the (a) length, (b) width, and (c) area distributions of microplastic particles found to contaminate blank controls. In (d) the counts of the different plastic and non-plastic particles present in each sample are shown. In (e) the composition of only the MPs detected in the different samples are shown. In (f) the number of MPs found in a human follicular fluid MPs sample (from Grechi et al. 2022) were corrected to the different blank controls presented in (e), and to the blank control that was ran in parallel with the follicular fluid sample (Fol. Fluid). Samples were corrected to the blanks by subtracting the number of particles detected in the water blanks from the corresponding number of particles detected in the follicular fluid sample for each specific plastic polymer individually.

Of the particles with HQIs ≥ 75, pigment related particles were the most abundant on average (mean ± SD = 36.3 ± 37.6 %), while non-plastic related particles accounted for 18.6 ± 13.8 % of the total number of identified particles, MP polymers for 9.4 ± 6.6 %, coating, solvents and fillers for 16.6 ± 17.7 %, and plasticizers for 4.7 ± 3.8 % (Fig. 1d). We also identified a total of 37 different MP polymers in the blanks, with silicon and polypropylene being the most abundant (mean ± SD = 27.8 ± 25.2 and 18.5 ± 14.4 %, respectively; Fig 1 e and table 1). The total concentration of MP particles varied substantially between samples, with a total of 23.50 particles mL^-1^ in laminar flow 1 (L. flow 1), 11.17 particles mL^-1^ in L. flow 2, 14.67 particles mL^-1^ in L. flow 3, 156.00 particles mL^-1^ in fume hood 1 (F. hood 1), 62.83 particles mL^-1^ in F. hood 2, 34.67 particles mL^-1^ in F. hood 3, 99.33 particles mL^-1^ in Bench 1, 9.50 particles mL^-1^ in Bench 2, and 29.67 particles mL^-1^ in Bench 3 (Fig. 1e; Table 1). A large number of fibres were also identified in all of the blanks. The lowest numbers of fibres were observed in blanks processed under laminar flow (mean ± SD = 27.8 ± 35.6 particles mL^-1^), followed by those processed on an open-air bench (56.1 ± 52.7 particles mL^-1^) samples, and lastly by blanks processed in a fume hood, which had an order of magnitude higher MP contamination (136.9 ± 107.9 particles mL^-1^).

**Table 1.** Different concentrations of particle polymers identified in the blank samples (particles/mL).

|  | Laminar flow |  |  | Fume Hood |  |  | Bench |  |  |
| --- | --- | --- | --- | --- | --- | --- | --- | --- | --- |
|  | 1 | 2 | 3 | 1 | 2 | 3 | 1 | 2 | 3 |
| ABS | 0.00 | 0.83 | 0.00 | 0.83 | 0.00 | 0.00 | 0.00 | 0.00 | 0.00 |
| CELLULOSE | 0.00 | 0.83 | 4.17 | 5.83 | 11.67 | 3.33 | 1.67 | 0.83 | 0.00 |
| NYLON | 0.00 | 0.00 | 0.00 | 0.00 | 0.00 | 0.00 | 2.50 | 0.83 | 0.00 |
| POLYPROPYLENE | 0.00 | 1.67 | 0.83 | 15.00 | 20.83 | 4.17 | 15.00 | 1.67 | 12.50 |
| POLYAMIDE | 0.00 | 0.00 | 0.00 | 1.67 | 0.00 | 0.00 | 0.00 | 0.00 | 0.83 |
| POLYESTER | 0.00 | 0.00 | 0.00 | 1.67 | 2.50 | 1.67 | 0.00 | 0.00 | 0.83 |
| POLYETHYLENE | 8.33 | 0.00 | 0.00 | 6.67 | 2.50 | 0.83 | 2.50 | 0.83 | 0.83 |
| POLYSTYRENE | 1.67 | 0.83 | 1.67 | 25.83 | 6.67 | 1.67 | 18.33 | 2.50 | 0.83 |
| POLYVINYL CHLORIDE | 0.00 | 0.00 | 0.00 | 1.67 | 0.00 | 0.00 | 0.83 | 0.00 | 0.00 |
| SILICON(IV) OXIDE | 12.50 | 2.50 | 0.00 | 77.50 | 5.00 | 17.50 | 50.00 | 0.00 | 0.83 |
| OTHER | 0.00 | 2.50 | 5.00 | 18.33 | 11.67 | 2.50 | 7.50 | 0.83 | 10.00 |
| **TOTAL** | **23.50** | **11.17** | **14.67** | **156.00** | **62.83** | **34.67** | **99.33** | **9.50** | **29.67** |

Our data demonstrate that using a fume hood results in the poorest quality blanks, with the highest amount of MPs and fibre contaminations, and also a high amount of variability between samples. Such contamination is expected, since fume hoods are designed to protect operators from hazardous substances by drawing laboratory air from outside of the hood to the inside of it. Using a laminar flow is the best method for reducing MPs contamination in the samples, as evidenced by the lowest amount of MPs detected in the blank controls. Our findings are in line with the results of a similar study by (Wesch et al., 2017), who investigated the contamination of filters placed in an indoor laboratory, in a mobile laboratory, in a fume hood and in a laminar flow cabinet by fibres. Fibres were detected in 95 and 89% of the filters from the indoor laboratory and the mobile laboratory (without any special air flow control), respectively. The lowest fibre detection was in the laminar flow filters (3.7%), while 50% of the fume hood samples presented microfiber contamination.

To demonstrate the importance of reliable, low-contamination blanks, we used raw data on the number of MPs detected in a human follicular fluid sample from Grechi et al. (2022) that was processed in a laminar flow. We then adjusted the MPs concentration in this sample to the 9 blanks we investigated here, as well as to the original blank from Grechi et al. (2022) that was produced in parallel to the sample processing. The blank correction was carried out by subtracting the number of particle detected in the water blanks from the corresponding particles detected in the follicular fluid sample, for each specific plastic polymer individually. As seen in figure 1f, the final number of MPs in the follicular fluid sample varied substantially across the different sets of blanks. Seven of the ten corrected samples included negative MPs polymer values, meaning that the blank control had more of the specific polymer than the original sample. If we were to have corrected the follicular fluid sample to the total number of MPs in either Bench 1 and Fume hoods 1 and 2 (*sensu* Liu et al., 2022; Tamminga et al., 2022), we might have concluded that no MPs were detected in the sample, since the total amount of MPs in these blank controls were higher than in the follicular fluid sample. Notably, even when the follicular fluid sample was corrected to the laminar flow blanks, there was still a large amount of variability in the final concentration of MPs. The high variability across blanks demonstrates how relying on only a single blank control can still generate inconsistent results, especially if samples are processed over multiple batches. We therefore recommend the use of separate blank controls processed under laminar flow in parallel with every sample collection, processing, and analysis process.

### 3.2 The use of blanks in the peer reviewed literature

#### 3.2.1 Nearly 20% of the current MPs research lacks controls against contamination

From the total 7,046 microplastics manuscripts identified on Pubmed, 2,869 were published in the past year and 718 had an Open Access license. We analyzed 59 of those 718 manuscripts, for their inclusion or not of blank controls and the conditions in which the samples were processed and analyzed. As shown in Data S3, around 83% of the manuscripts included blank controls in their experimental design. Of these, 92% provided the results from their blank controls, 26% did not detect any microplastics in their blanks, only 8% described the MPs size in their blanks, and 37% corrected the number of MPs detected in the blanks in their samples being analyzed.

#### 3.2.2 The majority of MP studies used ineffective blanks

Knowing that MPs are ubiquitous in the environment and a persistent source of contamination (Prata et al., 2020; see also Fig. 1), it was concerning to see that only 34% of the studies used a laminar flow cabinet, fume hood, clean room, or some other type of controlled air flow to process their samples. Further, almost 35% of the studies that described air control as a form of contamination prevention did so by using a fume hood, which was the least reliable approach of those we tested. In line with our empirical observations, 86% of the studies that used a fume hood reported contamination in their blanks. Unfortunately, however, none of them described the size ranges of the MP contaminants, so we could not compare these studies to our own fume hood blanks.

In addition to ineffective air flow control, the reported absence of MPs in the blank controls of 13 out of the 45 studies that showed their blank results was surprising given our experimental findings. We suspect that this was likely due to the size range of the particles analyzed in each of the studies, since MP contamination is normally caused by small particles that are airborne, attached to the surface of equipment, or in solutions (Prata et al., 2020); see also Fig. 2). A study from Frei et al. (2019), showed how most (62%) of the MPs detected in their blank controls were in the 20 to 50 μm size-range, while 30% of the particles were between 50 to 100 μm, 7% between 100 and 500 μm, and only less than 1 % were in the 500 to 1,000 μm range (Frei et al., 2019). Here, 7 out of 13 studies that did not detect microplastics in their sample controls included MPs < 50 μm and 6 investigated MPs bigger than 100 μm, suggesting that they might not have detected the majority of the plastic particles that contaminated the samples during processing (see also Fig. 2).

#### 3.2.3 Studies frequently prepare blanks, but do not use them to correct their data

Generating reliable information on the amount of MPs in an environmental or biological sample requires not only that we detect external contamination via blank controls, but also that we account for such contamination when analysing our samples. In this regard, of the 31 studies that detected MPs in their blanks, only 63% described how they corrected the amount of MPs detected in their samples to the amount of MP contamination in their blanks.

## 4. Conclusions

The past years have seen an incredible increase in the number of published articles about MPs. Yet, the growth in the number of MPs studies being published in recent years has not been accompanied by an increase in quality control of the studies performed. Thus, despite repeated calls, MPs research still lacks harmonization of the protocols used for collection, isolation and characterization of MPs that are critical for the future of the field (Ivleva, 2021; Provencher et al., 2020; Schymanski et al., 2021). Taken together with the lack of any standardisation in presenting the methods and data (Jenkins et al., 2022), reliably comparing and synthesising the results of MPs studies is an extremely challenging endeavour. Therefore, the field’s standards still need to improve substantially if we are to provide reliable information that can help address the plastic pollution crisis. Here we focused on only one of these overlooked methods for standardization: the use of adequate blank controls. We have shown that, due to the pervasiveness of MPs, the laboratory environment where samples are processed represents an important source of MPs contamination, which, in turn, challenges our ability to reliably detected MPs in different biological samples. Overall, properly planning, performing, and describing blank controls are imperative steps for ensuring the reliability and consistency of data and propel the field of microplastics research forward.

## Supporting information

PRISMA checklist

Table with all data from literature review

All detected particles

List of all detected particles and their classification

## CRediT authorship contribution statement

Conceptualization: MAMMF; Methodology: MAMMMF, NG, CLM, MJN; Investigation: NG and CLM; Supervision: MAMMF, MJN; Writing—original draft: MAMMF; Writing—review & editing: all authors.

## Supplementary data

Supplementary data S1: List of all detected particles and their classification into non-plastic related particle (NP), plastic polymer (PP), pigment (PG), plasticizer (PZ), coating, solvent and fillers (CSF), or fiber (FB).

Supplementary data S2: All detected plastics with hit quality index (HQI) ≥ 75 for all samples.

Supplementary data S3: Table with all data from literature review.

## Data availability

All data are available in the main text or as supplementary data.

## References

European Commission, 2018. A European Strategy for Plastics in a Circular Economy. Communication from the Commission to the European Parliament, the Council, the European Economic and Social Committee and the Committee of the Regions. Brussels.

Frei, S., Piehl, S., Gilfedder, B.S., Löder, M.G.J., Krutzke, J., Wilhelm, L., Laforsch, C., 2019. Occurence of microplastics in the hyporheic zone of rivers. Sci. Rep. 9, 15256. https://doi.org/10.1038/s41598-019-51741-5

Gardon, T., El Rakwe, M., Paul-Pont, I., Le Luyer, J., Thomas, L., Prado, E., Boukerma, K., Cassone, A.L., Quillien, V., Soyez, C., Costes, L., Crusot, M., Dreanno, C., Le Moullac, G., Huvet, A., 2021. Microplastics contamination in pearl-farming lagoons of French Polynesia. J. Hazard. Mater. 419. https://doi.org/10.1016/j.jhazmat.2021.126396

Grechi, N., Franko, R., Rajaraman, R., Stoeckl, J.B., Trapphoff, T., Dieterle, S., Frohlich, T., Noonan, M., de Almeida Monteiro Melo Ferraz, M., 2022. Microplastics are present in follicular fluid and compromise gamete function in vitro: Is the Anthropocene throw-away society throwing away fertility? bioRxiv 2022.11.04.514813. https://doi.org/10.1101/2022.11.04.514813

Hernandez, L.M., Xu, E.G., Larsson, H.C.E., Tahara, R., Maisuria, V.B., Tufenkji, N., 2019. Plastic Teabags Release Billions of Microparticles and Nanoparticles into Tea. Environ. Sci. Technol. 53, 12300–12310. https://doi.org/10.1021/acs.est.9b02540

Ivleva, N.P., 2021. Chemical Analysis of Microplastics and Nanoplastics: Challenges, Advanced Methods, and Perspectives. Chem. Rev. https://doi.org/10.1021/acs.chemrev.1c00178

Jenkins, T., Persaud, B.D., Cowger, W., Szigeti, K., Roche, D.G., Clary, E., Slowinski, S., Lei, B., Abeynayaka, A., Nyadjro, E.S., Maes, T., Thornton Hampton, L., Bergmann, M., Aherne, J., Mason, S.A., Honek, J.F., Rezanezhad, F., Lusher, A.L., Booth, A.M., Smith, R.D.L., Van Cappellen, P., 2022. Current State of Microplastic Pollution Research Data: Trends in Availability and Sources of Open Data. Front. Environ. Sci. 10. https://doi.org/10.3389/fenvs.2022.912107

Li, D., Shi, Y., Yang, L., Xiao, L., Kehoe, D.K., Gun’ko, Y.K., Boland, J.J., Wang, J.J., 2020. Microplastic release from the degradation of polypropylene feeding bottles during infant formula preparation. Nat. Food 1, 746–754. https://doi.org/10.1038/s43016-020-00171-y

Liu, Z., Bai, Y., Ma, T., Liu, X., Wei, H., Meng, H., Fu, Y., Ma, Z., Zhang, L., Zhao, J., 2022. Distribution and possible sources of atmospheric microplastic deposition in a valley basin city (Lanzhou, China). Ecotoxicol. Environ. Saf. 233, 113353. https://doi.org/10.1016/j.ecoenv.2022.113353

Munno, K., De Frond, H., O’Donnell, B., Rochman, C.M., 2020. Increasing the Accessibility for Characterizing Microplastics: Introducing New Application-Based and Spectral Libraries of Plastic Particles (SLoPP and SLoPP-E). Anal. Chem. 92, 2443–2451. https://doi.org/10.1021/acs.analchem.9b03626

Page, M.J., McKenzie, J.E., Bossuyt, P.M., Boutron, I., Hoffmann, T.C., Mulrow, C.D., Shamseer, L., Tetzlaff, J.M., Akl, E.A., Brennan, S.E., Chou, R., Glanville, J., Grimshaw, J.M., Hróbjartsson, A., Lalu, M.M., Li, T., Loder, E.W., Mayo-Wilson, E., McDonald, S., McGuinness, L.A., Stewart, L.A., Thomas, J., Tricco, A.C., Welch, V.A., Whiting, P., Moher, D., 2021. The PRISMA 2020 statement: An updated guideline for reporting systematic reviews. PLOS Med. 18, e1003583. https://doi.org/10.1371/journal.pmed.1003583

Paler, M.K.O., Leistenschneider, C., Migo, V., Burkhardt-Holm, P., 2021. Low microplastic abundance in Siganus spp. from the Tañon Strait, Central Philippines. Environ. Pollut. 284, 117166. https://doi.org/10.1016/j.envpol.2021.117166

Prata, J.C., Castro, J.L., da Costa, J.P., Duarte, A.C., Rocha-Santos, T., Cerqueira, M., 2020. The importance of contamination control in airborne fibers and microplastic sampling: Experiences from indoor and outdoor air sampling in Aveiro, Portugal. Mar. Pollut. Bull. 159, 111522. https://doi.org/10.1016/j.marpolbul.2020.111522

Provencher, J.F., Covernton, G.A., Moore, R.C., Horn, D.A., Conkle, J.L., Lusher, A.L., 2020. Proceed with caution: The need to raise the publication bar for microplastics research. Sci. Total Environ. 748, 141426. https://doi.org/10.1016/j.scitotenv.2020.141426

Ragusa, A., Svelato, A., Santacroce, C., Catalano, P., Notarstefano, V., Carnevali, O., Papa, F., Rongioletti, M.C.A., Baiocco, F., Draghi, S., D’Amore, E., Rinaldo, D., Matta, M., Giorgini, E., 2021. Plasticenta: First evidence of microplastics in human placenta. Environ. Int. 146, 106274. https://doi.org/10.1016/j.envint.2020.106274

Rillig, M.C., Kim, S.W., Schäffer, A., Sigmund, G., Groh, K.J., Wang, Z., 2022. About “Controls” in Pollution-Ecology Experiments in the Anthropocene. Environ. Sci. Technol. https://doi.org/10.1021/acs.est.2c05460

Schymanski, D., Oßmann, B.E., Benismail, N., Boukerma, K., Dallmann, G., von der Esch, E., Fischer, D., Fischer, F., Gilliland, D., Glas, K., Hofmann, T., Käppler, A., Lacorte, S., Marco, J., Rakwe, M. El, Weisser, J., Witzig, C., Zumbülte, N., Ivleva, N.P., 2021. Analysis of microplastics in drinking water and other clean water samples with micro-Raman and micro-infrared spectroscopy: minimum requirements and best practice guidelines. Anal. Bioanal. Chem. 413, 5969–5994. https://doi.org/10.1007/s00216-021-03498-y

Tamminga, M., Hengstmann, E., Deuke, A.-K., Fischer, E.K., 2022. Microplastic concentrations, characteristics, and fluxes in water bodies of the Tollense catchment, Germany, with regard to different sampling systems. Environ. Sci. Pollut. Res. 29, 11345–11358. https://doi.org/10.1007/s11356-021-16106-4

Tsering, T., Viitala, M., Hyvönen, M., Reinikainen, S.-P., Mänttäri, M., 2022. The assessment of particle selection and blank correction to enhance the analysis of microplastics with Raman microspectroscopy. Sci. Total Environ. 842, 156804. https://doi.org/10.1016/j.scitotenv.2022.156804

United Nations Environment Assembly, 2022. End plastic pollution: Towards an international legally binding instrument. Nairobi.

Wesch, C., Elert, A.M., Wörner, M., Braun, U., Klein, R., Paulus, M., 2017. Assuring quality in microplastic monitoring: About the value of clean-air devices as essentials for verified data. Sci. Rep. 7, 5424. https://doi.org/10.1038/s41598-017-05838-4

Xu, J.-L., Lin, X., Hugelier, S., Herrero-Langreo, A., Gowen, A.A., 2021. Spectral imaging for characterization and detection of plastic substances in branded teabags. J. Hazard. Mater. 418, 126328. https://doi.org/10.1016/j.jhazmat.2021.126328

